# Exome sequencing identifies variants associated with semen quality in Holstein Friesian and Hallikar bulls

**DOI:** 10.1101/2022.11.14.516500

**Authors:** Sarin K. Kunnath, K.P. Ramesha, Mukund A. Kataktalware, A. Kumaresan, S. Jeyakumar, D.N. Das, A. Manimaran, M. Joel Devadasan, A. Ashwitha, Shweta Mall, T.S. Keshava Prasad

## Abstract

Effective fertility of bulls is dependent on semen quality, often determined based on standard semen evaluation tests. Here we report Whole Exome Sequencing (WES) of 12 bulls from two breeds Holstien Friesian and Hallikar selected based on Ejaculate Rejection Rate (ERR). We explored the possibility of identifying genetic variants from the conserved protein coding regions of genome. A total of 10,510 SNPs and 10.236 INDELs were identified post alignment against reference genome (ARS-UCD 1.2) and were annotated using SnpEff. The number of variants with high and modifier functional impact detected were 145 and 19,122, respectively. Genetic variants common to both high and low ERR group bulls among Holstein Friesian were 08 and in Hallikarthe common variants were 51. Prominent *genesviz*. *UCP2, PANK2, GPD2, PTPRG, LARP7, EZH1, DENND1B* and *TDRD9* with a role in determining the semen quality were observed to be carriers of the genetic variant.

## Introduction

Improving the efficiency of reproductive ability of dairy cattle has been the biggest challenge for the dairy industry **[1]**. Selecting sires for improved fertility has been a task owing to its low heritability, compared to that of production traits **[2–3]** in cattle. While much attention had have been diverted on improving cow fertility traits, there requires a committed focus in understanding bull fertility traits **[1]**. The fertility of the breeding population either sire or dam determines the economic viability of livestock husbandry. More precisely sire fertility has greater impact than of the dam’s, as it is understood that sires are more than half the herd. Over the past few decades cattle breeds have been reared for improved milk production and beef, this preferential selection has resulted in development of breedswith specialised traits. The intensified preferential selection strategy has resulted in desirable production output, but with a consequent reduction in fertility traits owing to an antagonistic relationship between milk yield and fertility**[4]**. In the present circumstances focus should be on the need to upgrade the selection methodology with an aimto improve the fertility rate and produce viable off-springs to achieve intended target. This could probably be realised by including genetic information underlying traits such as semen functional and quality traits into selection strategy. Both candidate gene and genome-wide approaches have been successfully employed to study the genomic architecture of bull and have resulted in identifying bio-markers defining fertility **[1]**. A gene panel approach through genes associated with fertility and pathways regulating it, may be used as a filter to identify novel genetic variants. Hence, this study was planned with an aim to utilize whole exome sequencing, to unravel possible variants determining the bull semen quality in Hallikar a native cattle breed of Karnataka, India, known for its high endurance, hardiness and Holstein Friesian known for its high milk producing ability.

## Materials and Method

Holstein Friesian bulls maintained at Nandini Sperm Station, Hesarghatta, Bengaluru Karnataka, India and Hallikar bulls maintained at State Semen Collection Centre, Hesaraghatta, Bengaluru, Karnataka, India were used for the present study. Records on semen quality obtained for a period of one year from August 2019 to August 2020 were utilized to calculate Ejaculate Rejection Rate (ERR) using the formula: ERR (%) = Number of ejaculates found unsuitable for freezing ×100/ Total number of ejaculates obtained from the bull**[5]**. Based on ERR six (06) bulls each from Holstein Friesian and Hallikar breeds were selected and grouped as three (03) with lower ERR and three (03) with higher ERR **(S_1_)**. About 4 ml of blood from each selected bull were collected aseptically in a vacutainer tube containing 0.5% EDTA and gDNA was isolated using the Modified High Salt Extraction method. gDNA samples with 260/280 ratio ≥ 1.8 were utilised for the subsequent sequencing. Exome libraries were prepared by using Agilent SureSelect Bovine All Exon panel™ (kit) following the manufacturer’s protocol.

### Sequencing and Bio-informatic Analysis

Paired-end sequencing was carried out on IlluminaNovaseq 6000 platform, with a sequencing parameter of 150×2 at 100X depth. Data obtained through high throughput sequencing was subjected for a quality check using FastQC v0.11.9 **[6]** and base/reads with a Phred score (≥30) were opted and processed for base trimming, read filtering, adaptor clipping using fastp v0.20.0 **[7]**. The BWA v0.7.17 **[8]** index (parameters: -a bwtsw) and samtools were used to index the reference genome (GCF_002263795.1_ARS-UCD1.2) and also to support variant calling, respectively. Alignment of processed reads was carried out BWA mem v0.7.17 (parameters: ‘-M’). Variant call form prepared alignments were executed through GATK’s HaplotypeCaller (parameters: ERC GVCF) and GenotypeGVCFs tools. The filtered variants were thus scored using CNNScoreVariants (1D Model with pre-trained architecture) and annotated using snpEff v5.0d **[9]** to summarise the variants and predict their effects. Exonic intervals were obtained with Agilent SureSelect Bovine All Exon panels and were utilised in all supported steps of GATK and snpEff.

## Results

### Sequencing statistics

Whole Exome Sequencing (WES) of 12 bulls displayed a mean of 14.5 GB (8-19.2) **(S_2_)**. The mean percentage of duplication, GC content and alignment rate were 56.6, 48.3 and 99.99%. The mean number of bases with phred score ≥ Q30was 13532**(S_2_)**. Out of 20,746 genetic variants processed, 10510 SNP’s, 6765 Deletion and 3471 Insertionswere identified **(Fig1.)**. Number of genetic variants with high functional moderate and modifier impact identified are presented in **(Fig2.)**. The total number of transitions and transversions observed were 11115 and 6669 **(Fig3.)**, respectively. Shared and specific genetic variants observed between low and high ERR bulls in Holstein Friesian and Hallikar bulls are given in **(Fig4&5. S_3_)**.

**Fig 1.**
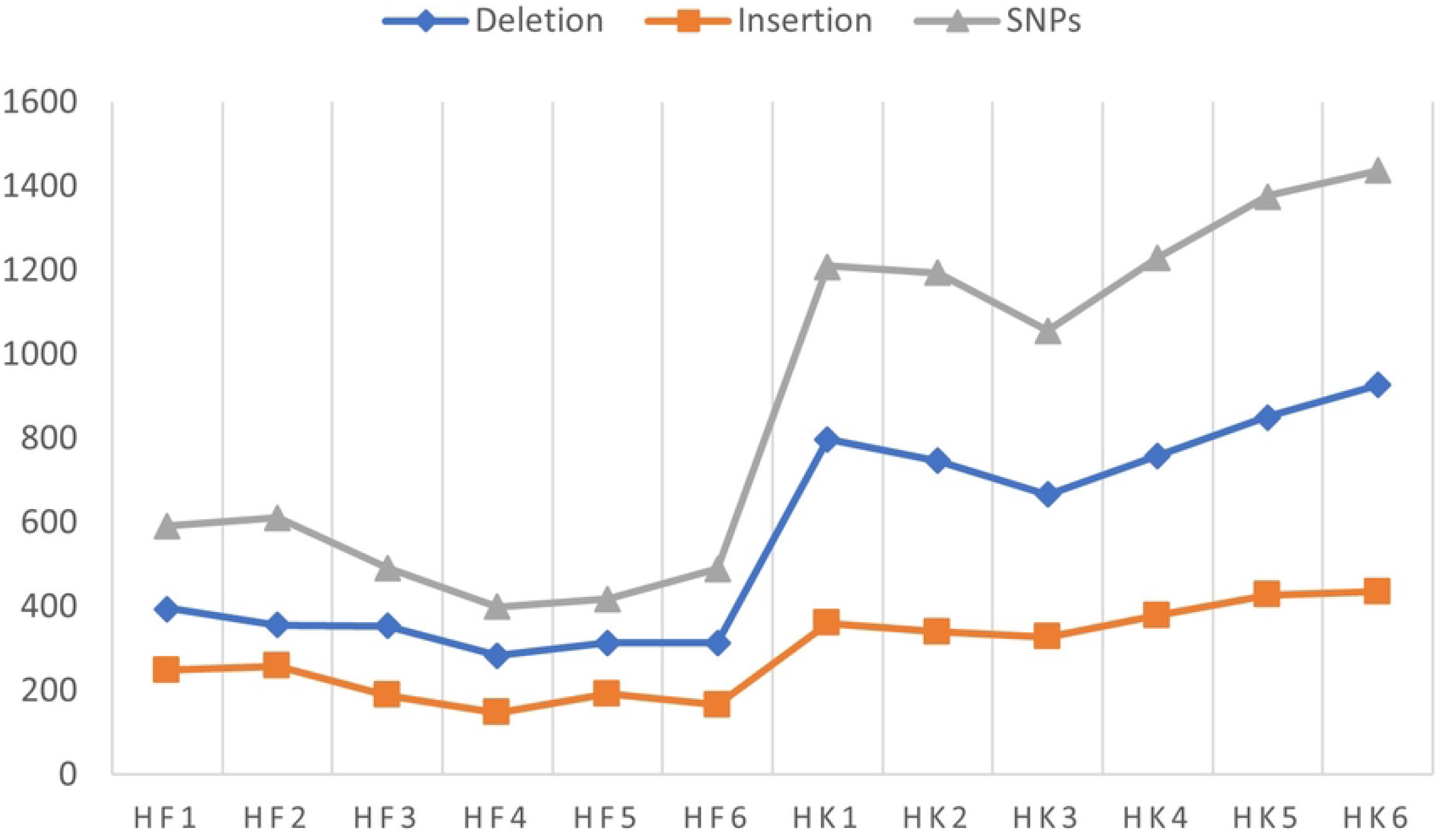
Genetic variants (*SNP, Insertions & Deletions*) identified through WES in Hallikar (HK) and Holstein Friesian (HF) Bulls.

**Fig 2.**
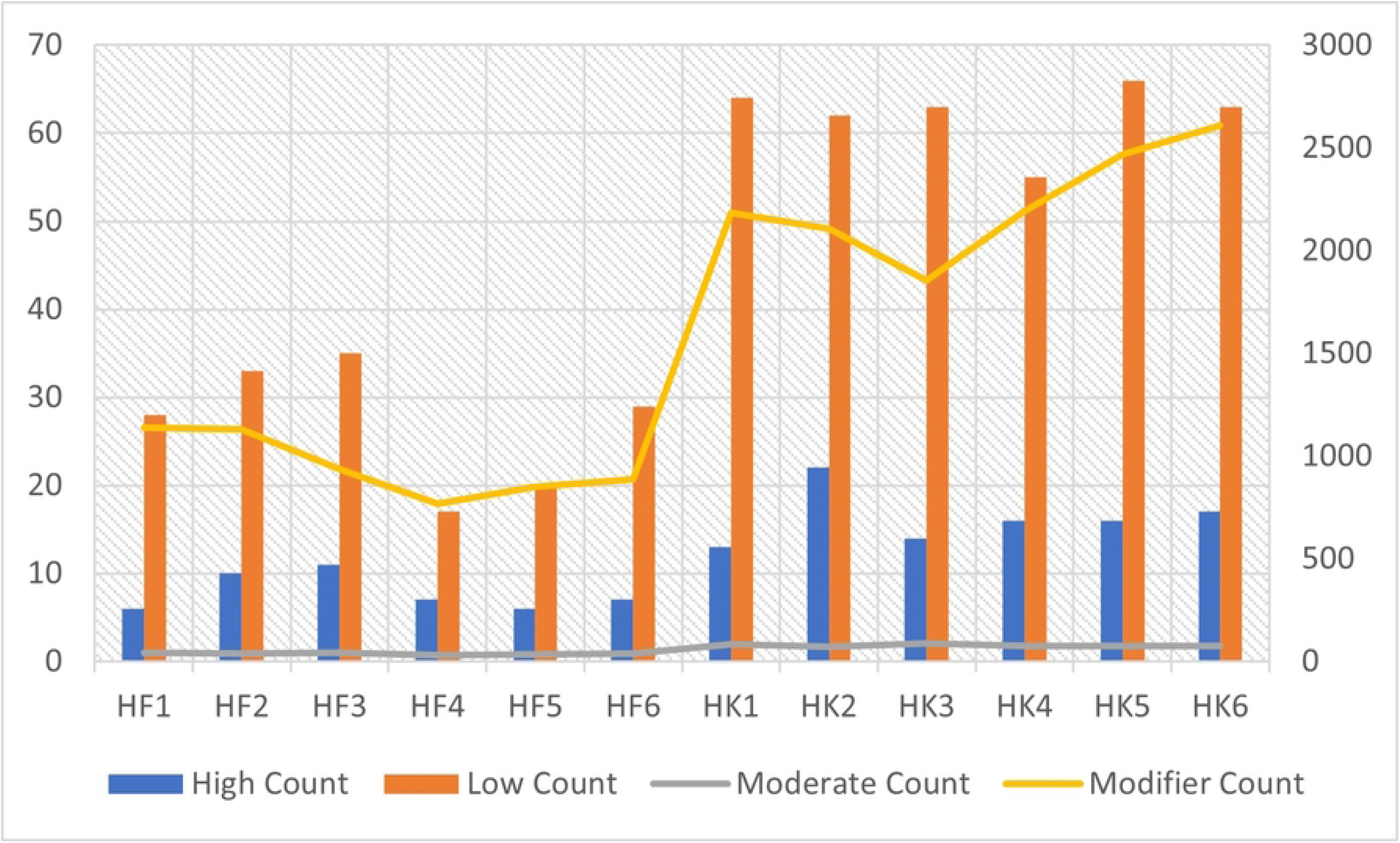
Effects of genetic variants (*High, Low, Moderate& Modifier*) identified through WES in Hallikar (HK) and Holstein Friesian (HF) Bulls.

**Fig 3.**
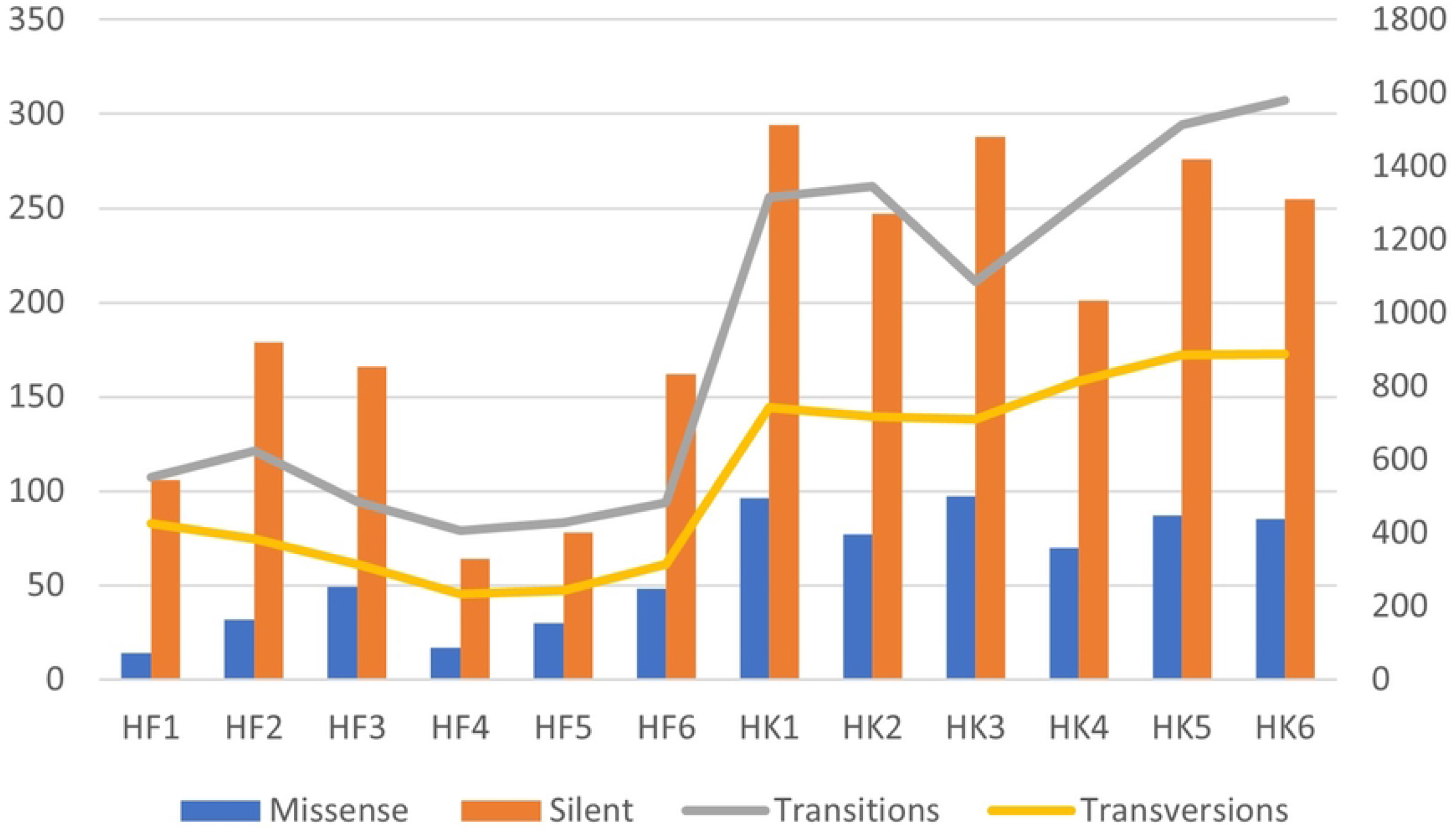
genetic variants by functional class (*Misense, Silent, Transitions & Transversions*) identified through WES in Hallikar (HK) and Holstein Friesian (HF) Bulls.

**Fig 4.**
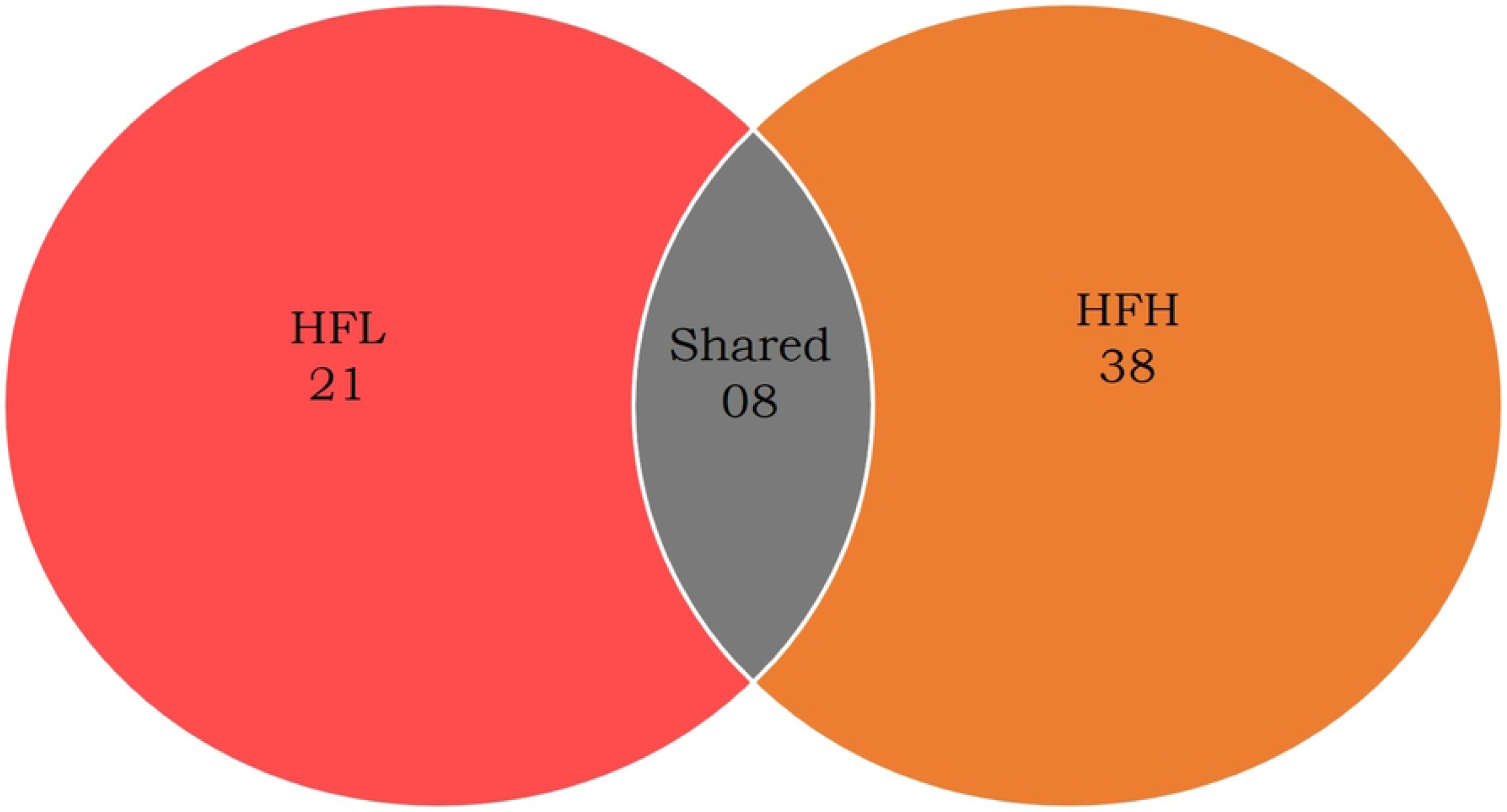
Venn Diagram Depicting Specific and Shared Variants (Semen Quality/Fertility Traits) in Holstein Friesian Low ERR (HFL) and High ERR (HFH) group Bulls.

**Fig 5.**
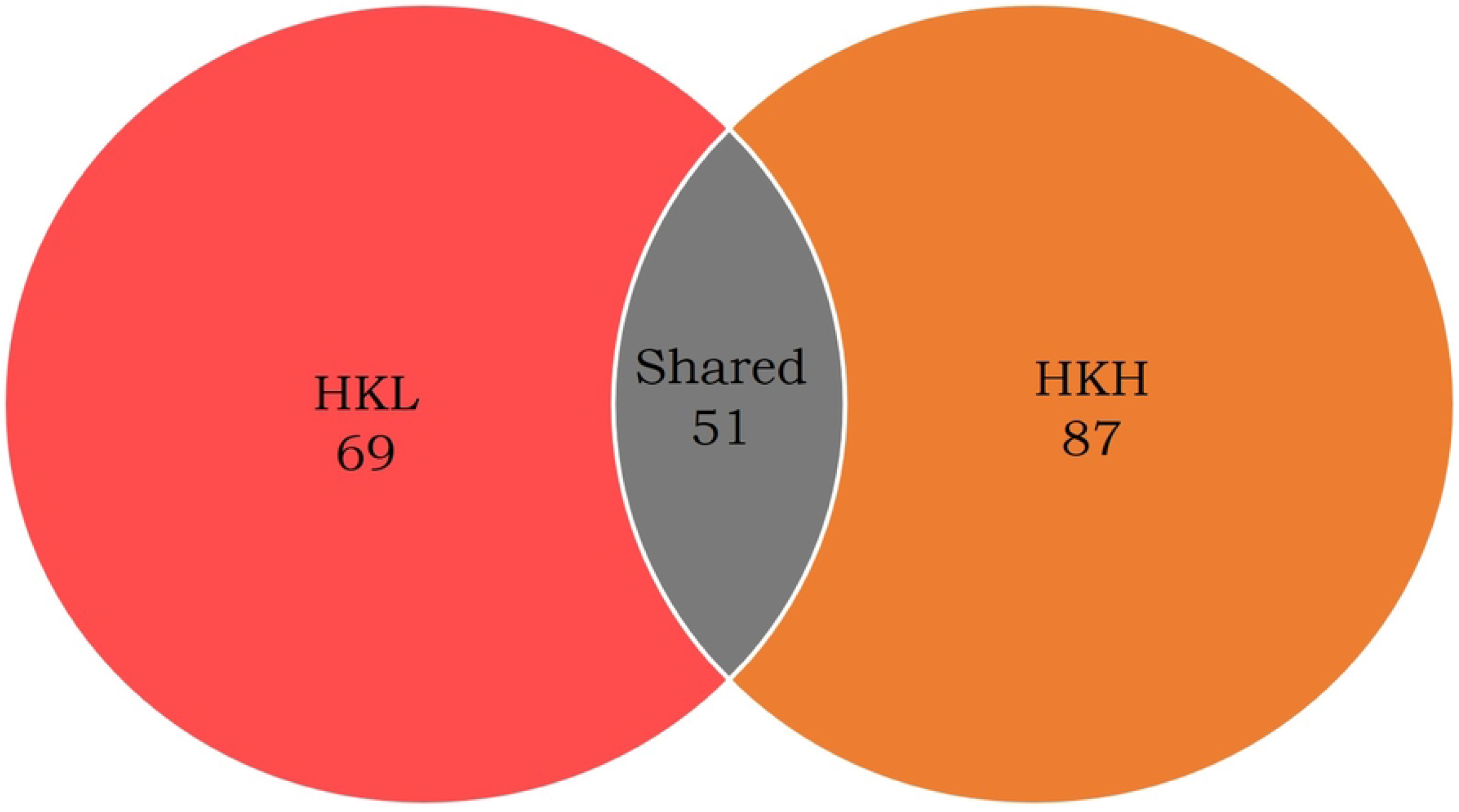
Venn Diagram Depicting Specific and Shared Variants (Semen Quality/FertilityTraits) in Hallikar Low ERR (HKL) and High ERR (HKH) Group Bulls.

## Discussion

Whole ExomeSequencing of 12 bulls resulted in identification of genetic variants**(S_4_)** among protein coding regions, which are concerned as the most conserved part of genome. The identified variants were lesser than those reported through WES carried out in Vechur cattle, Indian Buffalo breeds and Quarter horses [**10, 11 & 12]**.

The grouping of bulls was carried out on basis of ejaculate rejection rate. The minimal number of ejaculate rejections were grouped as low ejaculate rejection rate and those with maximum number of ejaculate rejections as high ejaculate rejection rate.Variants with previous reports suggesting relation with semen quality traits were identified from each group of bulls in both breeds.

We identified variants among genes *UCP2, TLE3, PAXIP1, CFAP6, DRD2, IFT140, TET, FER1l5, CCDC9, DNAJB13, TDRD* with previously reported relation to functional traits like, sperm motility, spermatogenesis, capacitation and sterility **[13–23]** in low ejaculate rejection rate group. Similarly in high ejaculate rejection rate group genes like *ADAM20, NPAS2, RYR3, DEFB124, TDRD9, TGFβ, SOX5* and *SOX6* related tosperm motility, capacitation, sterility, immunotolerance of sperm, maturation of spermatozoa immune defense system of the reproductive system **[24–30]** were observed. Few genetic variants among genes *likeADAM, AK7, ARID4B, CCDC9, CSMD1, DEFB124, DENND1B, DNAJB13, DRD2, FER1L5, PAX1p1, REC8, RYR3, TDRD9, TPCN1* and *WDFY* with previously reported relation to Male Fertility, Apoptosis, Immuno-tolerance, Sperm Motility, Capacitation and Spermatogenesis [**31–37**]were observed to be shared between both low and high ejaculate rejection rate bulls of both breeds under study.

### Annotation of genes identified in LowERR group

Among low ERR group bulls genes like *REC8, ACVR2A, PANK2, SOX4, LGR4, TBLIX1, WEE2, TDRD9, KDM3A and BRAC2* were identified to be involved in biological process pathways/functions**(S_5_)** concerned with determining the quality of semen. The important among them were spermatogenesis(GO:0007283), germ cell development (GO: 0007281), Wnt signalling pathway(GO:0016055), sex differentiation(GO:0007548), sexual reproduction (GO:0019953), reproductive process (GO:0022414), meiotic cell cycle (GO:0051321)and male gamete formation (GO: 0048232).

The lists of variants identified among genes with a role to play in semen functional traits were subjected to pathways analysis using DAVID database and the prominent among them were: Transforming growth factor-Beta signalling pathway, usually mediated through multiple pathways like canonical and non-canonical pathway, is a potent immune-modulator and its signalling is usually up regulated in disease conditions. Activin A and B **(38)** produced by gonads and Activins (ACVRA) are known to regulate spermatogenesis and steroidogenesis.

Actin Cytoskeleton (Pathway)plays an important role inproviding strength, connections to other cells and the extracellular matrix. Remodelling of cytoskeletal takes place during capacitation and involves actin polymerisation**(39)**. Through cyclic AMP/protein kinase pathway, fibronectin is known to stimulate capacitation of human sperm.

Vascular endothelial-derived growth factor (pathway)plays an important role in vasculogenesis and angiogenesis. Expression of VEGF in testes, prostate, seminal vesicles and semen have been studied and are believed to play crucial role in male fertility. The role of VEGF isoforms activity brings in significant changes in the ability of spermatogonial stem cell (SSC) to self-renew and colonize seminiferous tubules.

### Annotation of genes identified in High ERR group

The prominent functions/pathways**(S_6_)** observed among high ERR group bulls were Wnt signalling pathway (GO: 0016055), fertilization (GO: 0009566), gamete generation (GO: 0007276), sex differentiation (GO: 0007548), male meiosis (GO: 0007410), spermatogenesis (GO: 0007283) where in genes like *TLE3, TNFAIP3, DVL2, VCP, BRCA2, CD9, LRP6, IMMP2L, SIRT7, TXNDC8, OVGP1, REC8, KDM3A, LRP6* were observed to play a prominent role in regulating the pathways concerned with quality semen production.

The prominent pathways identified though pathway analysis using DAVID database were: Wnt signalling pathway, one of the most conserved pathway that regulates important aspects of cell fate determination, cell migration and organogenesis during embryonic development. Dishevelled protein through which biological signals are passed, are part of Planar Cell Polarity **(40)** and is known to regulate ectoplasmic specialization in testis through changes in cytoskeletal organization.

The primary regulator of mammalian reproductive function in both males and female is GnRH Biological Signalling Pathway and it acts via G-protein coupled receptors to stimulate synthesis and secretion of two gonadotropin hormones like luteinizing hormone and follicle-stimulating hormone **(41)**. Luteinizing hormone beta polypeptide (*LHβ*) gene is a single copy gene and it has been indicated that a mutation in the coding region of the gene could lead to its in activation and consequent impaired production of leydig cells **(42)** and leading to infertility in human. It also observed that the mutation would lead to susceptibly to prostate cancer via altered testosterone secretion.

Hypothalamicpituitary-testicular axis’s normal function is closely associated with male fertility **(43)**. Defective gene expression or abnormal signal transduction of hypothalamus and pituitary, have been observed to cause spermatogenic disorders and male infertility. PI3K-Akt Signalling Pathway are involved in multiple stages of male reproduction and have also been implicated in regulating of sperm autophagy, proliferation and differentiation of spermatogonia and somatic cells.

## Conclusion

The present study has resulted in identifying genetic variants in the conserved protein-coding regions of the genome. The variants identified were observed to be part of genes regulating major semen functional traits. Thus emphasising that Whole Exome Sequencing could be a viable option to identify genetic variants determining traits pertaining to reproduction and can be used as a tool in selecting bulls for breeding strategies.

## Acknowledgement

This study was supported and funded by the Ministry of Environment, Forest and Climate Change, Govt. of India under adaptation fund through Karnataka Livestock Development Agency (KLDA), Department of Animal Husbandry & Veterinary Services, Government of Karnataka under the project titled “Proteo-genomic approach to elucidate productive and reproductive performance of MalnadGidda, Hallikar, and Amrithmahal breeds of cattle and field performance recording of MalnadGidda cattle” to Southern Regional Station of ICAR-NDRI, Bengaluru. Sarin.K. Kunnath was deputed by P.V. Narsimha Rao Telangana Veterinary University for pursuing his doctoral programme at ICAR-NDRI;Joel.M. Devadasan and A. Aswitha were supported through ICAR-NDRI fellowship. The authors extend thanks to Molsys for carrying out Whole Exome Sequencing under out Sourcing. The authors acknowledge the support and assistance rendered by the Officers and staff of Nandini Sperm Station, Hesarghatta, Bengaluru Karnataka, India and State Semen Collection Centre, Hesaraghatta, Bengaluru, Karnataka, project staff of KLDA-MoEFCC funded project. The authors also record their sincere gratitude to the Director, ICAR-National Dairy Research Institute, Karnal for extending all the support and facilities for the project.

## Supporting Information

S_1_ Table 1. Holstein Friesian Bulls Selected Based on Ejaculate Rejection Rate (ERR).

Table 2.Hallikar (HK) Bulls Selected Based on Ejaculate Rejection Rate (ERR).

S_2_ Table 3.Whole Exome Sequencing Statistics of Raw Data in Holstein Friesian (HF) and Hallikar (HK) Bulls.

Table 4. General Statistics of Whole Exome Sequencing in Holstein Friesian (HF) and Hallikar (HK) Bulls.

S_3_ Table 5. Total Number of Shared Variants among Holstein Friesian (HF) and Hallikar (HK) Bulls.

S_4_ Table 6. List of identified genetic variants associated with semen and fertility traits group wise.

S_5_ Functional Annotation of genes (Genetic Variants) in Holstein Friesian (HF) and Hallikar (HK) Low ERR Group uisng DAVID database

S_6_ Functional Annotation of genes (Genetic Variants) in Holstein Frioesian (HF) and Hallikar (HK) High ERR Group uisng DAVID database

